# PERK inhibition mitigates restenosis and thrombosis - a potential low-thrombogenic anti-restenotic paradigm

**DOI:** 10.1101/581397

**Authors:** Bowen Wang, Mengxue Zhang, Go Urabe, Guojun Chen, Debra Wheeler, David Dornbos, Allyson Huttinger, Yitao Huang, Shahid Nimjee, Shaoqin Gong, Lian-Wang Guo, K Craig Kent

## Abstract

**Background:** Drug-eluting stents (DES) represent the main-stream management of restenosis following treatments of occlusive cardiovascular diseases. However, DES cannot eliminate in-stent restenosis yet exacerbate thrombogenic risks. To achieve dual inhibition of restenotic smooth muscle cell (SMC) de-differentiation/proliferation and thrombogenic endothelial cell (EC) dysfunction, a common target in both cell types, has been long-sought after. We evaluated the potential of protein kinase RNA-like endoplasmic reticulum kinase (PERK) as such a target for low-thrombogenic anti-restenotic intervention.

**Methods and Results:** We used a rat angioplasty model of restenosis and a FeCl_3_-induced mouse model of thrombosis. Loss-or gain-of-function was achieved by PERK inhibition (GSK2606414, siRNA) or overexpression (adenovirus). Restenosis was robustly mitigated by GSK2606414 administered either via injected (i.v.) lesion-homing platelet membrane-coated nanoclusters or a perivascular hydrogel; it was enhanced by PERK transgene. Whereas PERK inhibition blocked, its overexpression exacerbated PDGF-induced human aortic SMC de-differentiation (reduced smooth muscle α-actin or αSMA) and proliferation (BrdU incorporation). Further, PERK activity promoted STAT3 activation but inhibited SRF transcriptional (luciferase) activity; its protein co-immunoprecipitated with STAT3 and also MRTF-A, the SRF activator for αSMA transcription. Importantly, PERK inhibition also prevented TNFα-induced impairment of human EC growth and upregulation of thrombogenic tissue factor, both in vitro and ex vivo. In vivo, oral gavage of GSK2606414 preserved ~50% of the normal blood flow 60 min after FeCl_3_-induced vascular injury.

**Conclusions:** PERK inhibition is dual beneficial in mitigating restenosis and thrombosis, thus implicating a potential design for anti-restenotic intervention to overcome the thrombogenicity of DES.

## Introduction

Cardiovascular disease is the number one cause of death in developed countries. Angioplasty with stenting is the most commonly performed procedure to reconstruct occluded vessels, accounting for ~1 million cases worldwide and at a $11.18 billion cost in 2017 in the US alone. Unfortunately, reconstructed vessels, particularly those in the periphery, become re-narrowed; this recurrent disease is referred to as restenosis. Restenosis occurs primarily due to the formation of neointimal lesions in the vessel wall, a process termed intimal hyperplasia (IH). Its central etiology is smooth muscle cell (SMC) phenotypic switching and endothelial cell (EC) dysfunction[1, 2]. These two cell types are vital as they comprise the vasculature tunica media and endothelial inner lining.

Angioplasty mechanically damages SMCs and the endothelial lining, exposing SMCs and ECs to a myriad of stimuli such as growth factors and cytokines. In response, these two cell types undergo phenotypic changes and lose normal function. Specifically, phenotypically switched SMCs de-differentiate, becoming contraction-impaired and proliferative/migratory cells that form neointima[3, 4]. Meanwhile, dysfunctional ECs become growth-impaired and apoptotic, and produce thrombogenic factors (e.g. tissue factor) while releasing pro-inflammatory signal molecules that exacerbate SMC dysfunction and neointimal growth[5, 6]. Thus, an ideal anti-restenotic treatment should block not only SMC’s phenotypic switch but also EC dysfunction.

Drug-eluting stents (DES) are most commonly deployed to reduce post-angioplasty restenosis. However, they do not completely prevent IH and worsen thrombogenic risks[7]. In-stent restenosis occurs in ~10–23% of coronary DES applications, and >50% in peripheral arterial diseases. Moreover, stent-thrombosis has become a major concern; the most feared consequence is high rates (up to 40%) of sudden death of patients with stent thrombosis[8]. Increasing evidence reveals that DES mitigate the disease phenotypes of SMCs but not that of ECs; rather, they exacerbate EC dysfunction[9, 10] hence promoting IH and thrombosis. Therefore, the ultimate outcome of DES implantation for restenosis prevention, paradoxically, can be the persistence of in-stent restenosis and exacerbated thrombogenic risks in many patients[7]. Anticoagulant therapies cannot completely prevent stent thrombosis, and if applied long-term, can inflict significant financial cost and risk of bleeding[11]. To tackle the downsides of DES, alternative strategies such as drug-eluting balloons and biodegradable stents have been clinically tested. However, their applications still rely on EC-toxic drugs, and recent studies have brought to light increased mortality associated with their usage[12]. Therefore, there is a clear clinical need for an endothelium-protective and stent-free anti-restenotic therapy[13], where basic and translational research to address this need has been very limited in spite of recent progress[9, 14]. In order to rescue the endothelium thereby reducing thrombogenic risk while mitigating IH, it is critically important to identify and inactivate a common target in both SMCs and ECs that is responsible for their disease phenotypes.

Recently, endoplasmic reticulum (ER) stress response pathways have emerged as novel interventional targets for cardiovascular diseases. Three unique pathways, represented by inositol-requiring kinase 1 (IRE1), activating transcription factor 6 (ATF6), and protein kinase RNA-like endoplasmic reticulum kinase (PERK), are selectively or simultaneously activated depending on the ER stress context [15, 16]. Of note, the IRE1 and ATF6 pathways have been reported to play a significant role in restenosis and other vascular diseases[15, 16]. By contrast, the role of PERK in restenosis remained unclear. In this pathway, the ER-resident kinase PERK serves as the sensor for ER stress; its auto-phosphorylated form activates elongation initiation factor 2α (eIF2α). Activated eIF2α suppresses global protein translation while paradoxically activating the transcription of subsets of genes in a context-dependent manner, e.g. ATF4, a key counter-stress transcription factor[17]. In addition, activated PERK can also regulate other signaling pathways via its kinase activity[18]. Therefore, global translational repression, together with selective activation of a subset of genes as well as crosstalk with other pathways, all elicited by PERK activation, may define specific outcomes in a context (e.g. cell type) dependent manner[19].

Context dependence of PERK-mediated responses and implications from other two ER-stress response pathways led us to explore a potential involvement of PERK in restenosis. We found that PERK inhibition was robustly effective in stymying restenosis in a rat carotid artery angioplasty model. More significantly, we further uncovered that PERK inhibition was also anti-thrombotic in a FeCl_3_-induced mouse model. This finding was unexpected given that an anti-thrombotic effect has not been shown in the interventions of the other two pathways. Selective inhibitors for PERK have been well developed, and the first-in-class, GSK2606414, has demonstrated outstanding therapeutic efficacy in multiple disease models, including prion infection, Alzheimer’s disease, diabetes, hemorrhagic stroke, and cancers [20]. In the current study, we explored the potential of PERK inhibition as an anti-restenotic option featuring low thrombogenic risks, and the underlying cellular and molecular mechanisms.

## Methods and materials

### Ethics Statement

All animal studies conform to the Guide for the Care and Use of Laboratory Animals (National Institutes of Health publication No. 85–23, 1996 revision) and protocols approved by the Institutional Animal Care and Use Committee at the University of Wisconsin and the Ohio State University.

### Reagents and materials

Human aortic SMCs and ECs and their respective medium were purchased from Lonza (Walkersville, MD). BrdU ELISA colorimetric kit was purchased from Roche-Sigma Aldrich (Indianapolis, IN). Human recombinant PDGF-BB and TNF-α were purchased from R&D Systems (Minneapolis, MN). TRIzol, High Capacity cDNA synthesis kit, SYBR Green PCR Mastermix, and siRNAs (scrambled control and PERK-targeting siRNA) were purchased from Thermo Fisher (Waltham, MA). GSK2606414 was purchased from Apexbio (Houston, TX).

### Primary Vascular Smooth Muscle Cell and Endothelial Cell Cultures

Human aortic SMCs and ECs were cultured in SmGM-2 complete medium containing 5% FBS and EGM-2 complete medium containing 2% FBS, respectively. Cells between passage 5 and 7 were used for all experiments and maintained at 37 °C with 5% CO_2_. For cell culture expansion, 0.25% Trypsin was used for detachment of human SMCs, while Accutase (Thermo Fisher Scientific, Waltham, MA) was used for human ECs.

For siRNA-mediated PERK knockdown, small interfering ribonucleic acids (siRNAs) for PERK (*Eif2ak3*) were ordered and tested for efficiency. The validated siRNA (4390824, Silencer Select; Thermo Fisher Scientific, Waltham, MA) was transfected into SMCs by using RNAiMax Reagent (13778-075; Thermo Fisher Scientific, Waltham, MA) following the manufacturer’s protocol at a concentration of 50 nM.

For in vitro PERK overexpression, adenoviral vectors expressing GFP (Ad-GFP) and human PERK (Ad-PERK) were constructed using a one-step Adeno-X-ZsGreen adenoviral system (632267; Takara Bio, Mountain View, CA) following manufacturer’s instruction. Virus were amplified, purified and titrated as previously described[21]. SMCs were infected for 6 hr with adenoviruses (3×10^4^ particles/cell) in SmBm2 medium containing 2% FBS, and recovered for 20 h with 10% FBS, and then starved with 0.5% FBS for 24 h followed by treatments as illustrated in each experiment.

### Preparation of Delivery Tools for PERK Inhibitor In Vivo Application

Platelet membrane-coated nanoclusters were synthesized as previously described[13]. Briefly, PAMAM-PVL-OH was reacted with succinic anhydride with presence of 4-dimethylaminopyridine at a weight ratio of 100:16.4:25 to generate PAMAM-PVL-COOH for nanocluster synthesis. PAMAM-PVL-COOH was then dialyzed against distilled water to remove impurities, and then lyophilized for long-term storage. For nanocluster drug loading, PAMAM-PVL-COOH and PERK inhibitor GSK2606414 were dissolved in DMSO at a weight ratio of 5:2. Distilled water was then added drop wise into the above solution over the course of 30 min. The unloaded drugs and DMSO were subsequently removed by dialysis. Drug-loaded nanoclusters were then fused with platelet membrane vesicles under sonication for 2 min (100 μg of PAMAM to 20 μl of platelet membrane solution).

Perivascular tri-block gel was synthesized as previously described[22]. Briefly, heat-dried OH– PEG–OH was reacted with LA and GA at a weight ratio of 2.4:5:1.2 and then dried under vacuum for 30 min at 70 °C. Upon complete melting, the mixture was then added with Sn(Oct)_2_ as catalyst for initiation of the polymerization process, at a catalyst:[LA:GA] molar ratio of 1:500. After 8 hours of reaction at 150 °C, the mixture was dissolved in 4 °C cold water and then re-heated to 80 °C to precipitate the copolymers and remove other impurities; this process was repeated three times. The final product was lyophilized for long-term storage, and the triblock gel working solution was prepared by dissolving the polymer in water at 23% by weight.

### Cell Proliferation Assay

BrdU ELISA cell proliferation assay was performed following manufacturer instructions. For SMCs: SMCs were seeded in 96-well plates at a density of 4000 cells per well with a final volume of 200 μl, and starved for 24 h in SmBM-2 basal medium containing 0.5% FBS. Cells were then pretreated with 1 µM GSK2606414 or an equal volume of vehicle control (DMSO) for 2 hr in fresh starvation medium prior to PDGF-BB mitogenic stimulation (final 20 ng/ml). For ECs: ECs were seeded in 96-well plates at a density of 8000 cells per well with a final volume of 200 μl of EBM-2 medium containing 10% FBS. Cells were then pretreated with 1 µM GSK2606414 or an equal volume of vehicle control (DMSO) for 2 hr in fresh EBM-2 medium prior to TNF-α stimulation (final 20 ng/ml). 22 hr after stimulation, cells were labeled with BrdU by a 2 hr incubation at 37 °C, and then fixed with a FixDenat solution for 30 min, followed by a 90-min incubation at room temperature with an anti-BrdU-POD antibody (1:100 dilution). After washing with PBS for 3 times, substrate was added. Plates were incubated at room temperature for 30 min, and then colorimetric signals were measured on a FlexStation 3 Benchtop Multi-Mode Microplate Reader (Molecular Devices, Sunnyvale, CA) at 370 nm with a reference wavelength of 492 nm.

### Real-time Quantitative PCR (qRT-PCR)

mRNA was isolated from collected cells using TRIzol following the manufacturer's instructions. Purified mRNA (1 μg) was used for the first-strand cDNA synthesis and quantitative RT-PCR was performed using the 7500 Fast Real-Time PCR System (Applied Biosystems, Carlsbad, CA). Each cDNA template was amplified in triplicates using SYBR Green PCR Master Mix.

### Cellular and tissue protein extraction and Western blotting

SMCs and ECs were lysed in RIPA buffer containing protease inhibitors (50 mM Tris, 150 mM NaCl, 1% Nonidet P-40, 0.1% sodium dodecyl sulfate, and 10 μg/ml aprotinin). For tissue protein extraction, freshly harvested tissues were immediately snap-frozen in liquid nitrogen and completely pulverized with plastic mini-pestle in liquid nitrogen bath, followed by addition of RIPA buffer. Protein concentrations of cell or tissue lysates were determined using a Bio-Rad DC™ Protein Assay kit. Approximately 50 μg of proteins from each sample were separated on 4–20% Mini-PROTEAN TGX precast gels (Bio-Rad) and transferred to a PVDF membrane.

Proteins of interest were detected by immunoblotting using the following primary antibodies and dilution ratios: Rabbit anti-phospho-PERK (1:100) from Santa Cruz Biotechnology (sc-32577), rabbit anti-PERK (1:1000) from Cell Signaling Technology (3192), rabbit anti-phospho-eIF2α (1:1000) from Cell Signaling Technology (3398), mouse anti-phospho-STAT3 (1:1000) from Cell Signaling Technology (4113), rabbit anti-phospho-p65 (1:1000) from Cell Signaling Technology (3033), rabbit anti-MRTF-A (1:1000) from Cell Signaling Technology (14760), rabbit anti-CD142/Tissue Factor (1:500) from Thermo Fisher (PA5-27278), rabbit anti-CHOP (1:100) from Santa Cruz Biotechnology (sc-8327), rabbit anti-ATF4 (1:1000) from Abcam (ab184909), rabbit anti-Cyclin D1 (1:1000) from Abcam (ab134175), mouse anti-α–SMA (1:5000) from Sigma-Aldrich, and mouse anti-β-actin (1:5000) from Sigma-Aldrich. After incubation of the blots with HRP-conjugated secondary antibodies (1:3000 for goat anti-rabbit or 1:10,000 for goat anti-mouse, Bio-Rad), specific protein bands on the blots were visualized by applying enhanced chemiluminescence reagent Clarity Max Western ECL Substrate (Bio-Rad) and then recorded with a LAS-4000 Mini imager (GE, Piscataway NJ) and an Azure C600 imager (Azure Biosystems, Dublin, CA). Band intensity was quantified using ImageJ.

### Rat Carotid Artery Balloon Angioplasty

Carotid artery balloon angioplasty was performed in male Sprague–Dawley rats (Charles River; 300-350 g) as previously described[13]. Briefly, rats were anesthetized with isoflurane (5% for inducing and 2.5% for maintaining anesthesia). A longitudinal incision was made in the neck and carotid arteries were exposed. A 2-F balloon catheter (Edwards Lifesciences, Irvine, CA) was inserted through an arteriotomy on the left external carotid artery. To produce arterial injury, the balloon was inflated at a pressure of 1.5 atm and withdrawn to the carotid bifurcation and this action was repeated three times. The external carotid artery was then permanently ligated, and blood flow was resumed. Throughout the surgery, the animal was kept anesthetized via isoflurane inhaling at a flow rate of 2 L/minute. Carprofen (5 mg/kg) and bupivacaine (0.25% at incision site) were injected subcutaneously. Two weeks after balloon angioplasty, common carotid arteries were collected from anesthetized animals (under 2.5% isoflurane) following perfusion fixation at a physiological pressure of 100 mm Hg. Animals were euthanized in a chamber gradually filled with CO_2_.

In vivo overexpression of PERK in angioplastied carotid arteries was performed as we have previously described with modifications[23]. Briefly, immediately after angioplasty, an IV catheter (24GA, Insyte Autoguard, BD, Franklin Lakes, NJ) was inserted through the previously created arteriotomy on the external carotid artery and advanced past bifurcation while hemostats were still in place for common and internal carotid arteries. The catheter and external carotid artery were then tightly fastened using ligatures to prevent leaking. A syringe containing adenovirus solution was connected to the catheter, and its position was adjusted and secured using a flexible magnetic mount arm platform (Quadhands, Charleston, SC). A total of 150 ul adenoviral solution (>1×10^9^ IFU/ml) was then gradually infused into the common carotid artery to create local adenoviral infection, and incubated for 25 min. Saline-soaked gauge or Kimwipe were applied to keep the exposed carotid arteries and surrounding tissues from drying. Upon the completion of adenovirus solution infusion and incubation, the catheter was then withdrawn, and the common carotid artery lumen was then flushed repeatedly with saline containing 20U/ml heparin. Heparin was then injected (80U/kg, i.v.) to prevent thrombosis during the procedure, especially in the temporarily ligated internal carotid artery. Back flushing/bleeding from both common and internal carotid arteries was confirmed. In addition to Carprofen and bupivacaine, buprenorphine was also subcutaneously injected (0.05 mg/kg) for this particular experiment.

### Morphometric Analysis of Neointima

Paraffin sections (5 μm thick) were excised from carotid arteries at equally spaced intervals and then Verhoeff-van Gieson stained for morphometric analysis, as described in our previous reports[13]. Planimetric parameters as follows were measured on the sections and calculated using Image J: area inside external elastic lamina (EEL area), area inside internal elastic lamina (IEL area), lumen area, intima area (= IEL area – lumen area), and media area (= EEL area – IEL area). Intimal hyperplasia was quantified as a ratio of intima area versus media area.

Measurements were performed by a researcher blinded to the experimental conditions using 3– 6 sections from each of animal. The data from all sections were pooled to generate the mean for each animal. The means from all the animals in each treatment group were then averaged, and the standard error of the mean (SEM) was calculated.

### Luciferase assay for SRF transcriptional activity

Luciferase assay was performed as previously described[24]. Briefly, human primary aortic SMCs were transfected with pGL4.34 Vector plasmids (E1350; Promega, Madison, WI) using Effectene Transfection Reagent (301425; Qiagen, Germantown, MD). After screening with Hygromycin B, SMCs were then recovered for another passage and then seeded in 24-well plates at a density of 20,000 cells/well in complete SmGm2 medium after detached using Accutase. Cells were then infected with adenovirus as aforementioned and cultured for additional 24 hr before lysis in Bright-Glo (Promega; catalog no. 2610). Luminescence signals were measured on a FlexStation 3 Benchtop Multi-Mode Microplate Reader (Molecular Devices, Sunnyvale, CA), and CellTiter-Glo luminescence readings from duplicate wells were used for normalization.

### Co-immunoprecipitation Assay

All co-immunoprecipitation assays were performed in MOVAS SMC cell lines (CRL-2797; ATCC, Manassas, VA) as previously described[24]. Briefly, cells were stably transfected with either HA or HA-STAT3 vectors, followed by selection with ampicillin and subsequent treatment with PDGF-BB and GSK2606414. At the end of each treatment, cells were collected by using an immunoprecipitation lysis buffer (87787; Thermo Fisher Scientific, Waltham, MA) containing a protease inhibitor cocktail (78430; Thermo Fisher Scientific, Waltham, MA), and kept on ice for 30 min to ensure complete lysis. Cell lysates of equal protein amounts were incubated overnight at 4°C (constant rotation) with 2 μg of an anti-HA antibody or immunoglobulin G (IgG) control (sc2027; Santa Cruz Biotechnology, Dallas, TX). Co-immunoprecipitation of STAT3 with associated proteins was then performed by using Protein A/G Agarose beads included in the Pierce Classic IP Kit (26146; Thermo Fisher Scientific, Waltham, MA) followed by immunoblotting to detect immunoprecipitated HA/HA-STAT3 or co-immunoprecipitated PERK and MRTF-A.

### FeCl_3_-induced Arterial Thrombosis Model

Adult C57BL/6J wild type male and female mice were administered either GSK2606414 at the dosage of 150 mg/kg or vehicle control as a single oral gavage. Compound was mixed in 0.5% carboxymethyl cellulose and 0.05% Tween. 4 hours after treatment, mice were anesthetized with ketamine (55 mg/kg) plus xylazine (15 mg/kg) and rectal temperatures were maintained at 37°C by a thermo-regulated heating pad (Physitemp TCAT-2DF Controller Clifton, NJ). Non-invasive blood pressure by tail cuff measurement was recorded at baseline before injury and 60 minutes after injury. (CODA Kent Scientific Torrington, CT). The common carotid artery was gently dissected and a pulsed Doppler flow probe (MA0.5PSB; Transonic Systems; Ithaca, NY) placed to record blood flow velocity. Electrocardiography leads were placed to allow monitoring of ECG in addition to continuous heart rate recording throughout the entire procedure (LabChart, ADInstruments, Sydney, AU). Carotid artery injury was induced by application of Whatman filter paper (1 mm^2^) saturated with 10% FeCl_3_ solution on the adventitial surface proximal to the flow probe for 3 minutes. The flow as a percentage of baseline and the time to thrombotic occlusion (blood flow, 0 mL/minute) was measured from the placement of the FeCl_3_-saturated filter paper. Animals were reanesthetized as needed with ketamine to maintain deep anesthesia throughout the procedure and sacrificed 60 minutes after initiation of FeCl_3_ application. Both FeCl_3_-applied carotid arteries and the contralateral uninjured carotid arteries were collected for histology purposes.

### Ex vivo culture of Aortic Rings

Male Sprague–Dawley rats (Charles River; ~400 g) were sacrificed and perfused with cold heparinized Hanks balanced salt solution (HBSS). Thoracic aortas were collected, with all branches carefully dissected and removed in heparinized HBSS. Aortas were cut into rings of ~5 mm in length, and cultured in DMEM medium supplemented with 1% FBS in 24-well plates. Rings were evenly distributed in each well, and were pre-treated with 1 μM GSK2606414 at least 2 hr prior to stimulation with TNF-α at a final concentration of 100 ng/ml. 4 hr post stimulation, aortic rings were rinsed and snap-frozen in liquid nitrogen for protein extraction.

### Statistical Analysis

Data are presented as mean ± standard error of the mean (SEM). Statistical analysis was conducted using either Mann-Whitney nonparametric test or One-way ANOVA followed by post hoc bonferroni test, as specifically stated in figure legends. Data are considered statistically significant when a P value is < 0.05.

## Results

### Endovascular Delivery of PERK Inhibitor via a Biomimetic Nano Approach Markedly Mitigates Neointima Lesion and Restenosis in a Rat Balloon Angioplasty Model

Recently, we have developed a novel biomimetic platform that could achieve stent-free targeted delivery to vascular lesions[13]. We set out to apply this technology to the preclinical therapeutic evaluation of PERK inhibition in the context of anti-restenotic intervention, as a concerted effort to provide a safer and effective alternative to DES. In our previous study, platelet membrane-coated (biomimetic) nanoclusters proved to be non-immunogenic and able to home to the injured artery wall following i.v. injection in the rat balloon angioplasty model[13]. To evaluate the effect of PERK inhibition on IH, we tail-vein injected these biomimetic nanoclusters loaded with GSK2606414 (dosage of 2.5 mg/kg) immediately after balloon angioplasty. The second injection followed at day 5 post angioplasty (Figure 1A). At day 14, arteries were collected for histological analysis, which showed a profound anti-restenotic effect of GSK2606414 compared to vehicle control, with a >70% reduction in the intima to media (I/M) ratio (Figure 1C) and ~30% increase in lumen area (Figure 1D). Of note, this 2.5 mg/kg dose is ~1/5 of that used in a peri-vascular hydrogel delivery route (see below, Figure 2A). More significantly, it is only about 1/800 of that in a 14-day delivery regimen (150 mg/kg, b.i.d, po) commonly used in the literature[20, 25]. To the best of our knowledge, we have provided herein the first in vivo evidence showing the anti-restenotic efficacy of PERK inhibition. In addition, delivery of PERK inhibitor via an injectable biomimetic nanoplatform may cater to the clinical need for stent-free anti-restenotic therapies.

**Fig. 1.**
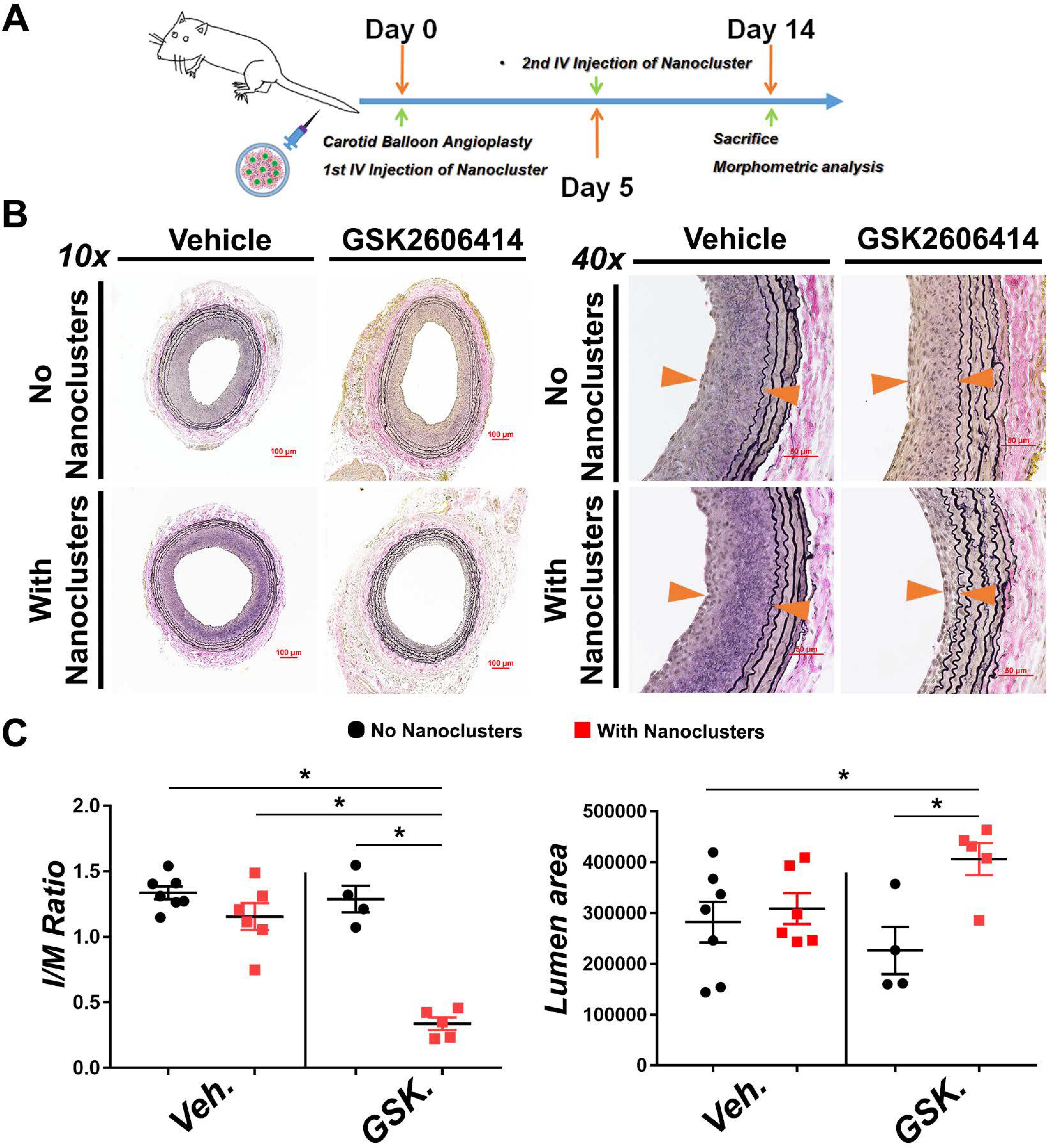
Endovascular targeted delivery of PERK inhibitor via a biomimetic nano approach mitigates IH in the rat model of carotid artery angioplasty. Drug delivery via biomimetic (platelet membrane-coated) nanoclusters that home to balloon-injured sites of rat carotid arteries was established in our recent report[13]. We used this targeted delivery method in the same rat angioplasty model for administration of vehicle (DMSO control) and PERK inhibitor (GSK2606414, 2.5 mg/kg). Biomimetic nanoclusters were tail-vein injected twice (immediately and 96h after angioplasty). Arteries were collected 14 days after angioplasty for morphometric analysis. A. Schematic illustration of tail-vein administration of PERK inhibitor carried in biomimetic nanoclusters. B. Representative Verhoeff-van Gieson (VVG)-stained sections. Neointima lesion is indicated between arrow heads. Scale bar: 100 μm for 10x, 50 μm for 40x. C. Quantification of IH (the intima/media area ratio) and lumen area. Mean ± SEM, n ≥4 animals; *p<0.05, One-way ANOVA with Bonferroni post hoc test.

**Fig. 2.**
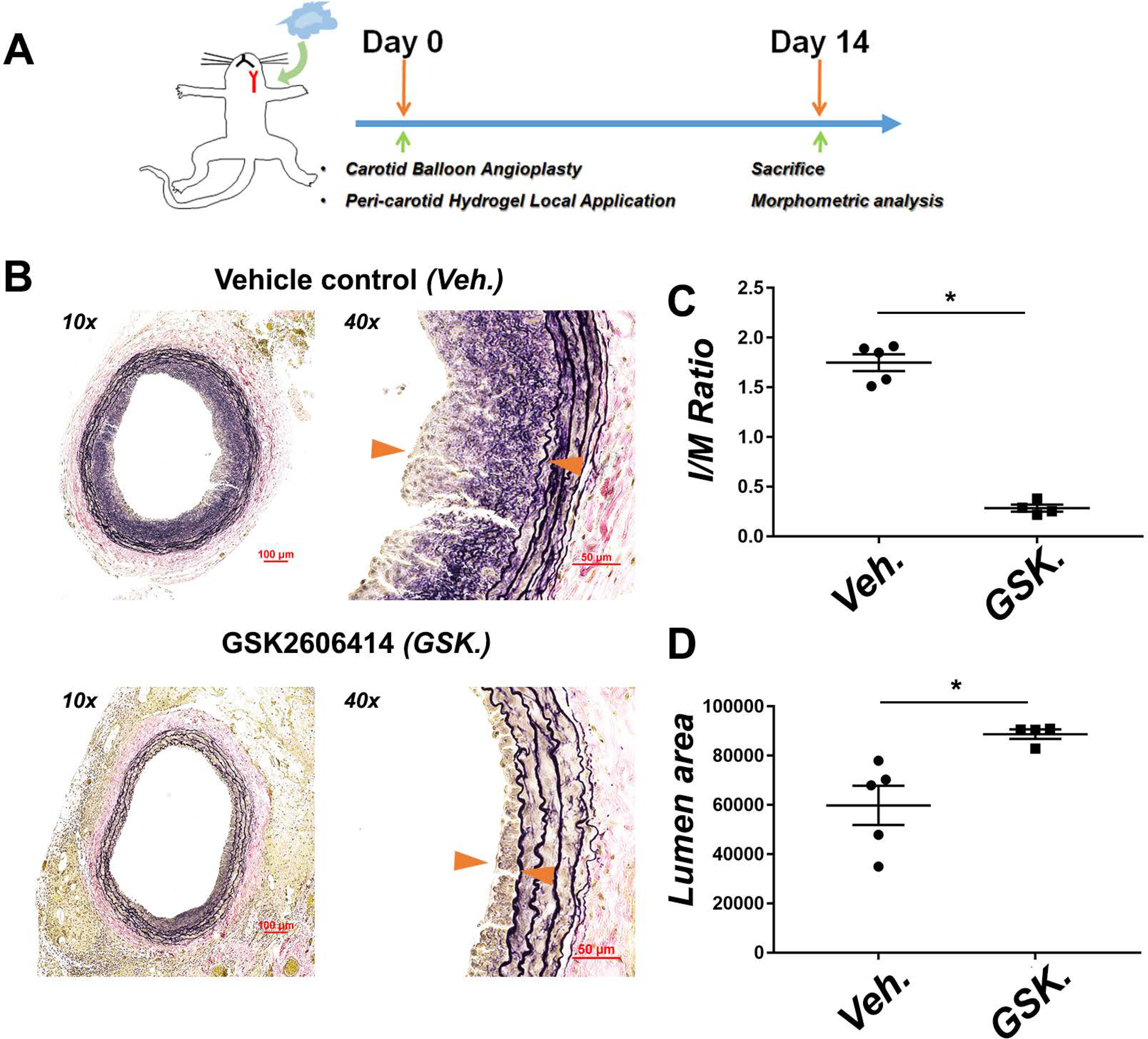
Perivascular local delivery of PERK inhibitor mitigates IH. Vehicle or PERK inhibitor (GSK2606414, 25 mg/kg) carried in a slow-release tri-block gel was perivascularly applied immediately after balloon angioplasty. Arteries were collected 14 days after angioplasty for morphometric analysis. A. Schematic illustration of periadventitial (common carotid artery) administration of PERK inhibitor carried in a triblock hydrogel. B. Representative VVG-stained sections. Neointima lesion is indicated between arrow heads. Scale bar: 100 μm for 10x, 50 μm for 40x. C. Quantification of intima/media (I/M) area ratio. D. Quantification of lumen area. Mean ± SEM, n= 4 rats; *p<0.05, Mann-Whitney nonparametric test.

### Perivascular Local Delivery of PERK inhibitor Effectively Ameliorates Neointima Lesion and Restenosis

To draw a solid conclusion on the anti-restenotic efficacy of PERK inhibition and to broaden its potential therapeutic application, we next tested GSK2606414 in a perivascular delivery route in the same rat balloon angioplasty model. To this end, we used a PLGA-PEG-PLGA thermosensitive triblock gel (liquid on ice and semisolid at body temperature), which in our previous studies exhibited excellent stability and a steady sustained (~4-week) drug release profile[22]. Using this gel system we applied GSK2606414 (25 mg/kg, one-time delivery) to the peri-adventitial space immediately after angioplasty. Morphometric analysis of arteries collected at day 14 indicated an ~80% reduction of the I/M ratio (Figure 2C) and an ~50% increase of lumen area (Figure 2D) in comparison to vehicle control. These results confirmed a robust anti-restenotic effect of the PERK inhibitor. Moreover, since this perivascular delivery method is in line with a prospective anti-restenotic management of open surgeries (e.g. bypass vein grafts)[26], PERK inhibition has a potential to be broadly applied.

### The PERK Pathway is Activated in Balloon-injured Arteries; its Overexpression Increases Neointima

Given the anti-restenotic efficacy of PERK inhibitor, we next determined a possible injury-induced activation of the PERK pathway. Uninjured and injured rat common carotid arteries at day 3, day 7, and day 14 post angioplasty were harvested and their homogenates were used for Western blot analysis of PERK pathway proteins. The data showed that dramatic PERK activation (a 4-fold increase of phospho-PERK) occurred in the artery 3 days after injury and persisted throughout the 14-day time course (Figure 3A and 3B). In accordance, the direct substrate of PERK and a central effector kinase, eIF2α, was also activated at day 3 (Figure 3A and Figure S1A). The decline of its activation at later time points suggests its acute response to the early-phase injury to control excessive ER stress[27]. ATF4 as an eIF2α downstream transcription factor is another central player in the PERK pathway. Its expression showed a pattern very similar to that of PERK activation (Figure 3A and Figure S1B). Considering that ER stress activates eIF2α which suppresses global translation yet allows for upregulation of a set of “privileged” genes[28], the temporal discrepancy in eIF2α activation and ATF4 upregulation possibly reflects a selective activation of ATF4’s expression. CHOP is a downstream transcription factor of the extended PERK pathway and is often associated with cell death when cellular damage is too severe[29]. Its upregulation at day 3 (Figure 3A and Figure S1C) coincided with the well-known peak of apoptosis frequently observed in this rat model[30]. Collectively, these results indicate that the PERK pathway was strongly activated due to arterial injury, consistent with a role in neointima development and its targetability by an inhibitor as evidenced above (Figures 1 and 2).

**Fig. 3.**
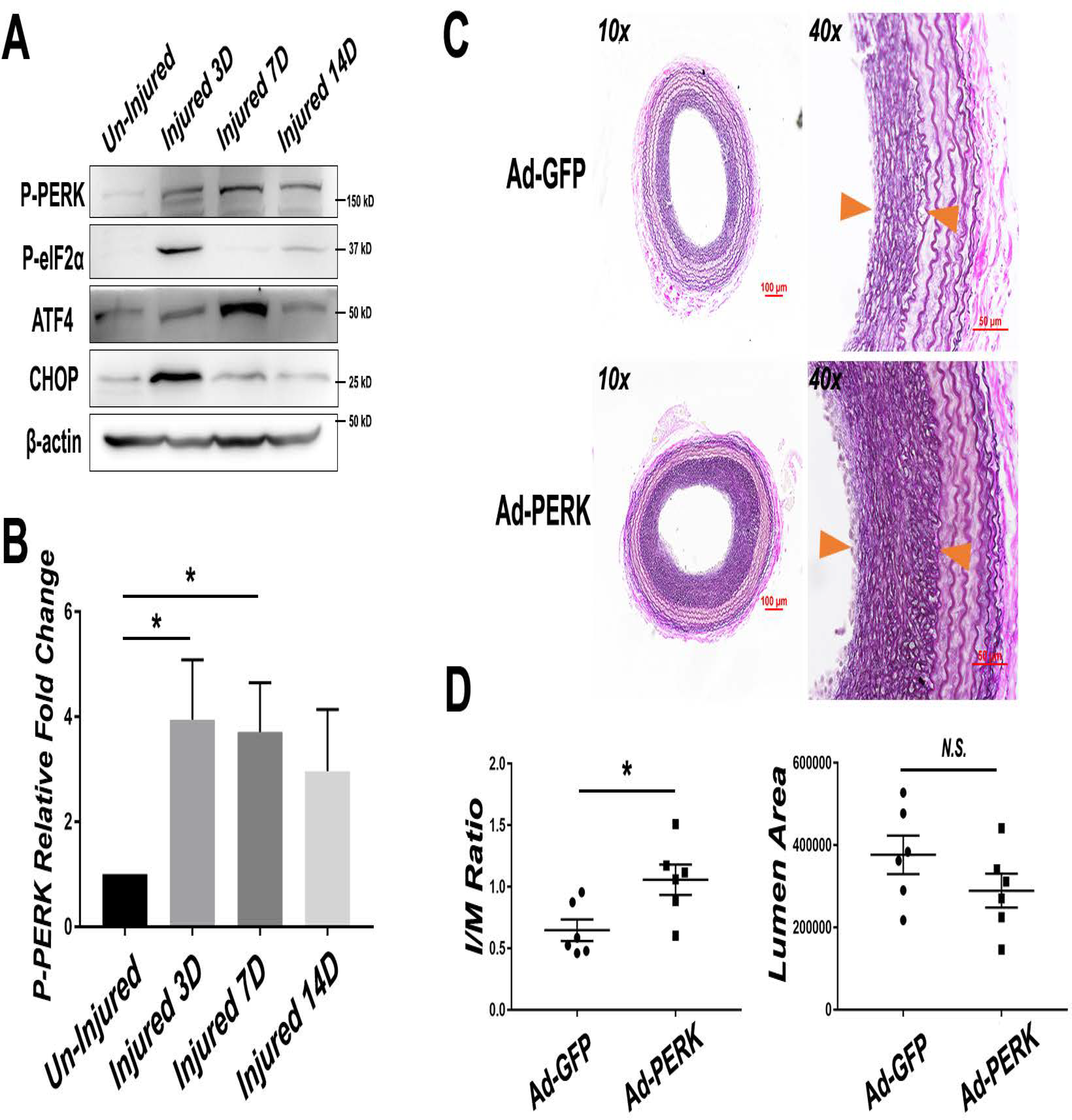
Injury-induced activation of the PERK pathway and exacerbated IH in balloon-injured rat carotid arteries overexpressing PERK. A. Activation of the PERK pathway in injured arteries. Following angioplasty, rat common carotid arteries were harvested at days 3, 7, and 14 (3d-14d) and homogenized for immunoblotting analysis. B. Quantitation of PERK phosphorylation level following rat balloon angioplasty. Mean ± SEM, n=3 rats; *p<0.05, One-way ANOVA with Bonferroni post hoc test. C. In vivo PERK gain-of-function. Ad-GFP or Ad-PERK was intraluminally infused with the injured artery for 25 min immediately after balloon angioplasty. Shown are representative VVG-stained sections. Neointima lesion is indicated between arrow heads. Scale bar: 100 μm for 10x, 50 μm for 40x. D. Quantification of intima/media (I/M) area ratio and lumen area. Mean ±SEM, n= 6 rats; *p<0.05, Mann-Whitney nonparametric test.

We then further verified PERK’s specific role in promoting neointima, via its overexpression using adenovirus intraluminally infused into the artery wall following the endothelium-denuding angioplasty. Effective transduction of PERK-expressing adenovirus was confirmed by Western blot analysis of artery homogenates (Figure S2A). Morphometric analysis of post-angioplasty 14-day artery sections indicates a significantly higher I/M ratio in arteries transduced with PERK-expressing adenovirus compared to the Adeno-GFP empty vector control (Figure 3C and 3D). Additionally, using tissue mRNA samples from a portion of the injured carotid artery, we also observed hallmarks of phenotypic switching, including reduced expression of α-SMA (Figure S2B). Therefore, these gain-of-function results support a specific role for PERK in promoting neointima formation.

### PERK Activation Promotes SMC Phenotype Switching

Phenotypic switching of SMCs is the core cellular mechanism underlying the development of neointima lesion and hence restenosis[4]. We therefore determined the specific function of PERK in this process via loss- and gain-of-function experiments. We used PDGF-BB, a widely used potent inducer, to stimulate SMC phenotypic switching which was measured as enhanced de-differentiation (decrease of α-SMA) and proliferation (BrdU incorporation and CDK1 expression). As shown in Figure 4A, whereas treatment of human primary aortic SMCs with PDGF-BB stimulated PERK activation (Figure 4A, 4E, and 4I) and cell proliferation (Figure 4C, 4G, and 4K) and reduced α-SMA by 2-3 fold (Figure 4D, 4H, and 4L), pretreatment with the PERK inhibitor GSK2606414 (1 μM) abrogated these PDGF-induced changes (Figure 4, A-D). We then confirmed this result via gene knockdown using a PERK-specific siRNA. The data indicate that PERK silencing *vs* scrambled siRNA control closely phenocopied the effects of GSK2606414 on SMC phenotypic switching (Figure 4, E-H). Importantly, consistent evidence was also derived from the gain-of-function experiments. As shown in Figure 4I-4L, PDGF-BB stimulated SMC proliferation and CDK1 expression and reduced α-SMA, this effect was enhanced in cells transduced with PERK-expressing adenovirus compared to empty-vector control. Combined, these in vitro loss- and gain-of-function results support a specific role of PERK in promoting SMC phenotypic switching.

**Fig. 4.**
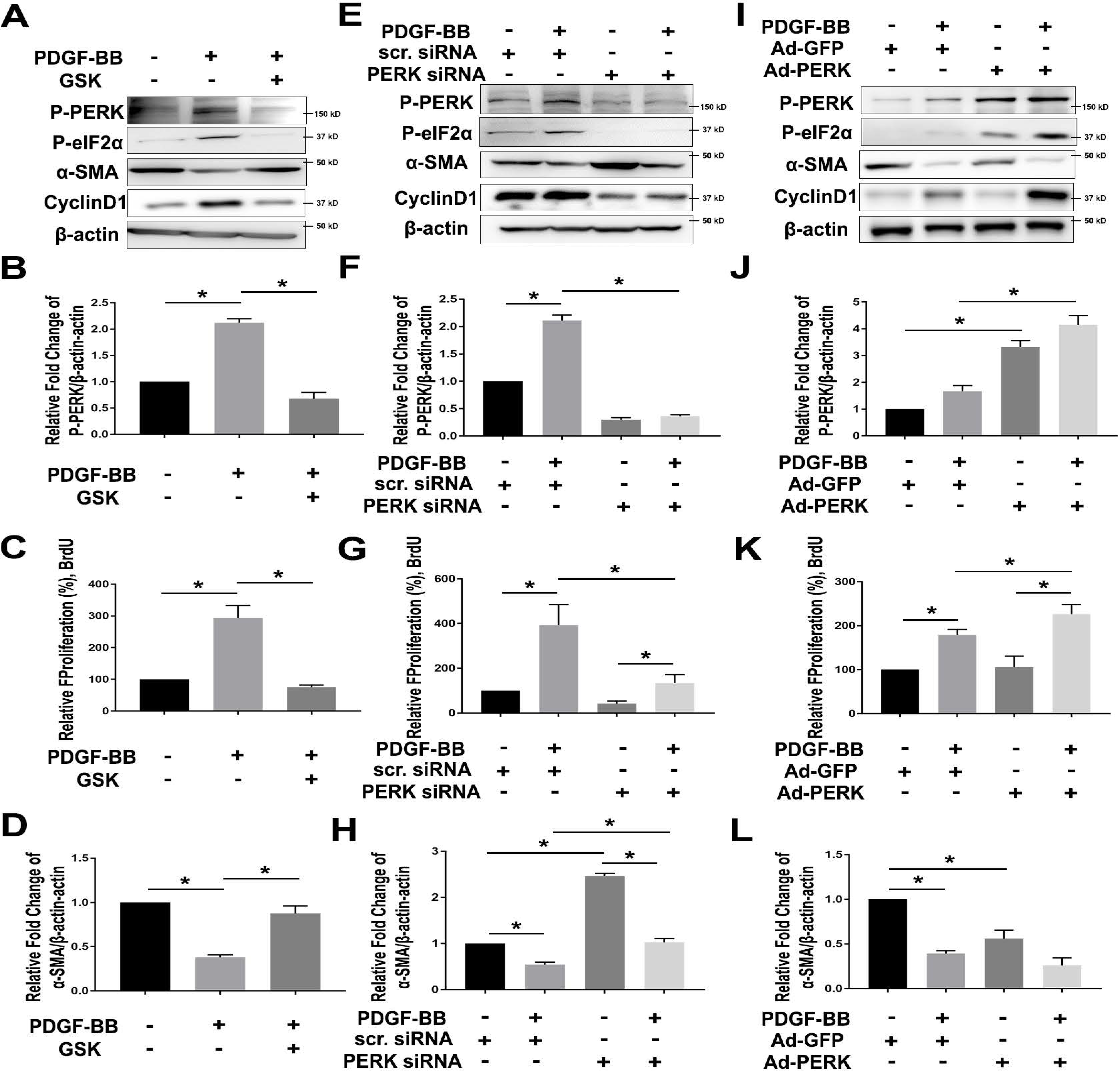
PERK loss- and gain-of-function respectively abrogates and exacerbates SMC phenotypic switching. To determine the specific role of PERK in SMC phenotypic switching (reduced α-SMA and increased proliferation), human primary aortic SMCs were cultured, starved, and pretreated with PERK inhibitor (A-D), siRNA (E-H), or adenovirus (I-L), and then treated with solvent or PDGF-BB (100 ng/ml). Cells were harvested 24 hours after PDGF stimulation for Western blot analysis of PERK pathway proteins and BrdU proliferation assay. A-D. Pharmacological blockade of PERK kinase activity. Cells were pre-treated with vehicle (DMSO) or 1 μM GSK2606414 for 2 hours before adding PDGF-BB. A: Representative immunoblots of PERK pathway (P-PERK and P-eIF2α) and SMC phenotypic switching (αSMA and CyclinD1). B: Proliferation of SMCs were determined using BrdU incorporation ELISA assay; n=3. C and D: Quantitation of PERK phosphorylation level and αSMA expression in SMCs stimulated with PDGF-BB and PERK inhibitor pretreatment. Protein band densitometry was normalized to β-actin; n=3. E-H. PERK genetic silencing. Starved SMCs were transfected with 50 nM PERK-specific siRNA for 48 hours, and then starved for 24 hours before PDGF stimulation. Similar to A-D, immunoblots of PERK pathway and SMC phenotypic switching markers are presented (E), together with proliferation assay (F; n=4) and quantitation of PERK phosphorylation level (G; n=3) and αSMA expression (H; n=3). I-L. Adenovirus-mediated PERK gain-of-function. Cells were transduced with Ad-GFP or Ad-PERK for 6 hours, recovered for 24 hours and then starved for 24 hours prior to PDGF stimulation. Similar to A-D and E-H, immunoblots of PERK pathway and SMC phenotypic switching markers are presented (I), together with proliferation assay (j; n=4) and quantitation of PERK phosphorylation level (K; n=3) and αSMA expression (L; n=3). All data are presented as mean ± SEM, *P < 0.05, one-way ANOVA with Bonferroni post hoc test.

### PERK Activation is Paralleled by STAT3 Activation and SRF Inhibition upon SMC Phenotypic Switching

To investigate the molecular mechanism underlying PERK’s functional role in SMC phenotypic switching, we explored its possible regulation of the determinants of SMC phenotypes. SRF is a well-established master transcription factor that governs αSMA expression and SMC differentiation; myocardin family proteins such as MRTF-A act as co-factors by activating SRF[31]. It is well-documented that suppressed MRTF/SRF signaling leads to SMC phenotypic switching[32]. In addition, recent studies have shown that STAT3 is highly sensitive to a range of cytokines, and it acts as a major mediator of SMC phenotypic switching by binding myocardin family proteins hence sequestering them from activating SRF[33]. Interestingly, we observed that PERK inhibition with either GSK2606414 or siRNA nearly abolished PDGF-induced STAT3 activation (Figure 5A and 5B). Further, co-IP experiments showed that HA-STAT3 specifically pulled down endogenous PERK in a PDGF-dependent manner, and this STAT3/PERK association was blocked by pretreatment with the PERK inhibitor (Figure 5C). Moreover, PDGF-induced and PERK inhibitor-blocked co-IP was also observed between HA-STAT3 and MRTF-A, similar to the literature evidence[34]. We were thus prompted to determine whether PERK regulates the SRF transcriptional activity. Indeed, PERK overexpression (*vs* GFP control) markedly inhibited the SRF activity in SMCs transfected with a luciferase vector containing the SRF-binding motif (CArG box) (Figure 5D).

**Fig. 5.**
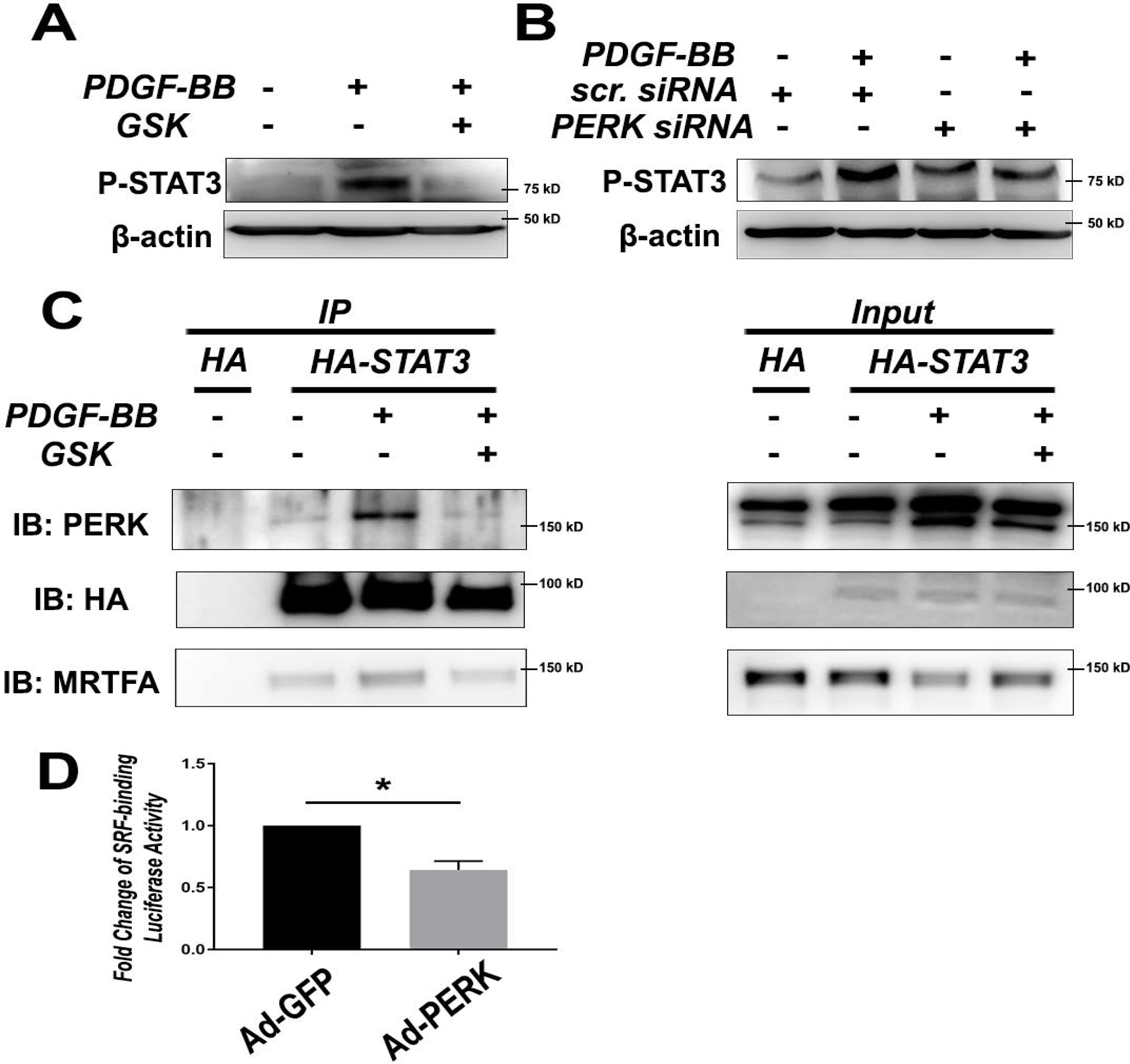
PERK regulates the activation of transcription factors STAT3 and SRF. SMCs were cultured, starved, and pretreated with PERK inhibitor (A) or siRNA (B) and then treated with solvent or PDGF-BB (100 ng/ml) for 24 hours as described previously. A and B. Immunoblots of STAT3 phosphorylation in SMCs. Shown are representative blots from two similar experiments. C. Co-immunoprecipitation. HA-STAT3 lentivirus or HA vector control was transfected into MOVAS prior to pre-treatment with PERK inhibitor and treatment with PDGF-BB. Cell lysates were enriched using HA antibody, and then probed for PERK and MRTFA. D. Luciferase assay for SRF transcriptional activity. Human aortic SMCs were transfected with a vector containing SRF response elements. Putative SRF activity was compared between two conditions: adenoviral overexpression of PERK (Ad-PERK) or GFP control (Ad-GFP), n=5; Mean ± SEM, *P < 0.05, Wilcoxon nonparametric test.

Up to this point, our data revealed a novel role for PERK in exacerbating neointima formation in injured arteries, and coherently, in mediating PDGF-induced SMC phenotypic switching. This PERK function in SMCs was at least partially accounted for by its positive regulation of STAT3 activation that has been previously implicated as inhibitory for the MRTF/SRF axis[33].

### PERK Inhibition Rescues TNFα-induced Human Primary Aortic EC Dysfunction

Bearing in mind the EC-toxic defect of DES – the *status quo* management of restenosis – we were motivated to evaluate the impact of the PERK-targeting strategy on EC health. ECs are extremely susceptible to inflammatory challenges; a cytokine storm following endovascular injury is known as notoriously detrimental to the homeostasis of endothelium. We used TNFα as a cytokine-storm mimic to induce EC dysfunction, as broadly performed in previous studies[35]. EC dysfunction manifests mainly as growth impairment and elevated production and secretion of tissue factor, which is a potent pro-inflammatory factor and thrombosis initiator[9, 10]. As shown in Figure 6, treatment of human primary aortic ECs with TNFα activated PERK and dramatically upregulated tissue factor by 40 fold (Figure 6A) and hampered EC proliferation (BrdU assay, Figure 6D); pretreatment with the PERK inhibitor almost completely averted these changes. Importantly, TNFα-induced upregulation of PERK and tissue factor as well as its abrogation by PERK inhibitor was replicated ex vivo using rat aortic artery explants (Figure 6F). At the transcription level, NFκB is known as a master transcription factor that dictates EC homeostasis or dysfunction if overactivated in a cytokine environment[36]. Consistently, we observed TNFα-stimulated p65 activation, which was attenuated by PERK inhibition (Figure 6E). Taken together, these results support a positive role for PERK in EC dysfunction in addition to that in SMC phenotypic switching. This finding is significant as PERK-mediated EC dysfunction (in particular tissue factor upregulation) has not been previously reported. Moreover, restoration of EC homeostasis via PERK inhibition implicates a potential approach to lowering thrombogenic risks.

**Fig. 6.**
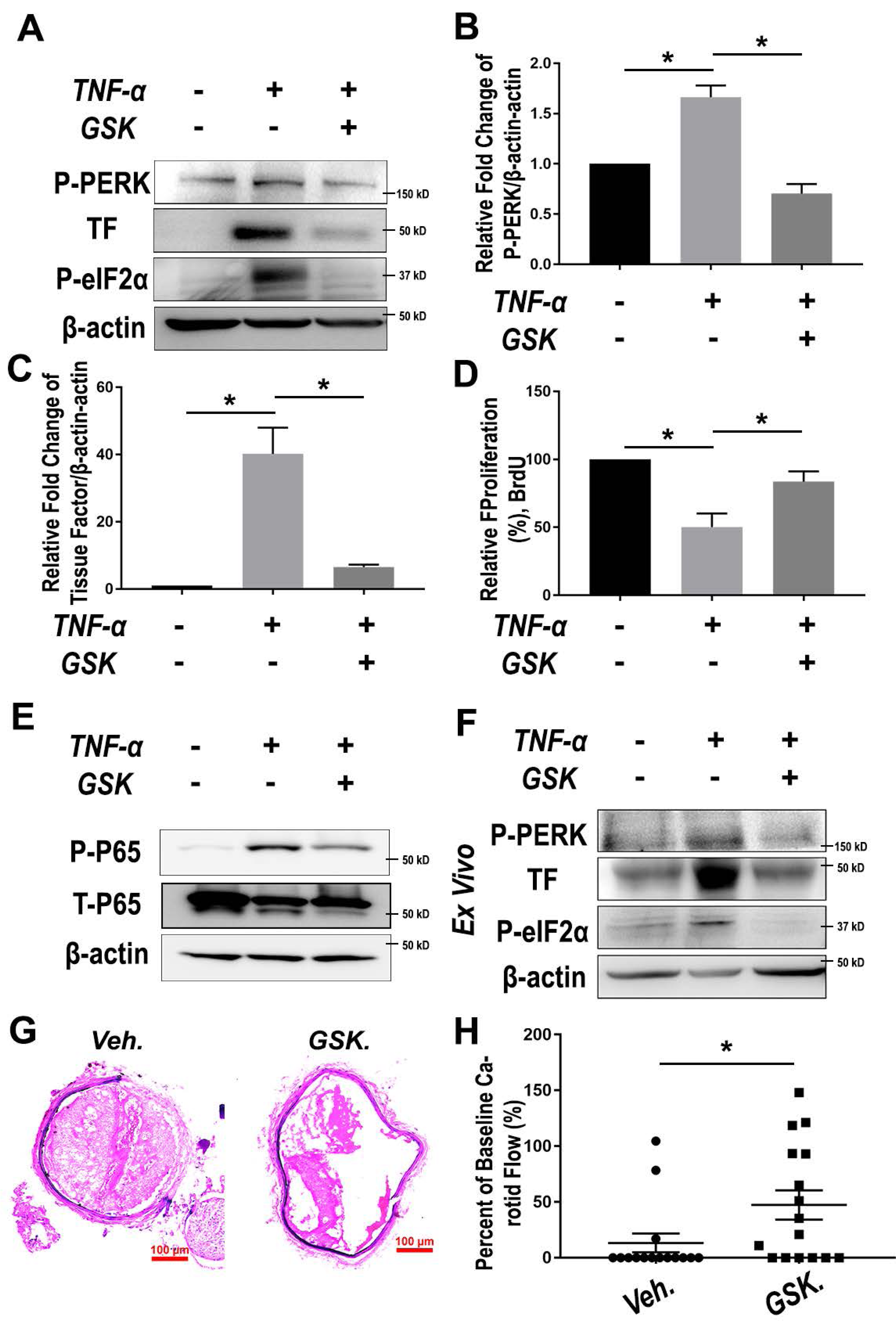
PERK Inhibition rescues EC dysfunction and mitigates thrombosis. A-E. Human primary aortic ECs were pre-treated with vehicle or 1 μM GSK2606414 for 2h prior to TNF-α stimulation (20 ng/ml). Cells were harvested at 6 hours and 24 hours after TNF-α treatment for immunoblotting and BrdU assays, respectively. p65 phosphorylation was determined 15 min after TNF-α stimulation. A: Representative immunoblots of PERK pathway (P-PERK and P-eIF2α) and EC dysfunction marker (thrombogenic marker Tissue Factor/TF). B and C: Quantitation of PERK phosphorylation level and TF expression in ECs stimulated with PDGF-BB and PERK inhibitor pretreatment. Protein band densitometry was normalized to β-actin; n=3. D: Proliferation of ECs was determined using BrdU incorporation ELISA assay; n=3. Data in B-D are presented as mean ± SEM, *P < 0.05, one-way ANOVA with Bonferroni post hoc test. E: Representative immunoblots of NFκB pathway (p65 phosphorylation). Shown are representative blots from two similar experiments.?? F. Ex vivo PERK pathway activation by TNF-α and its blockade by pretreatment with GSK2606414. Cultured rat aortic rings were pre-treated with vehicle or 1 μM GSK2606414 for 2 hours prior to a 6-hour stimulation with TNF-α (20 ng/ml). Shown are representative blots from two similar experiments. F-G, Mice were pre-treated with either vehicle or GSK2606414 (150 mg/kg) via oral gavage for 4 hours, and then subject to FeCl_3_topical application to induce carotid artery thrombosis. F: Representative H&E staining of FeCl_3_-injured carotid arteries from mice treated with vehicle or GSK2606414. G: Scatter plots of arterial flow at the end of the 60 min recording. A 0% flow indicates complete occlusion from thrombosis. Mean ± SEM, n = 15-16 mice. *P < 0.05, Mann-Whitney nonparametric test.

### Systemic Application of PERK Inhibitor Mitigates Thrombosis in a FeCl_3_-induced Murine Model

Inspired by effective repression of tissue factor in vitro and ex vivo via PERK inhibition, we finally tested an anti-thrombotic strategy in vivo. We used a widely recognized FeCl_3_-induced thrombosis mouse model that has been established in our previous studies[37]. GSK2606414 was administered via oral gavage 4h prior to FeCl_3_ application. Based on the literature information regarding the pharmacokinetics and pharmacodynamics of GSK2606414 and its derivatives, we chose a regimen of 150 mg/kg, which was estimated to result in an effective local level at the endothelium. After treating the animals with the PERK inhibitor, we topically applied FeCl_3_ around the carotid artery to initiate acute endothelium damage and a coagulation signaling cascade. Interestingly, despite a slight decrease in carotid artery flow in the first 20 minutes, pre-treatment of the mice with GSK2606414 maintained a higher arterial perfusion (Figure 6G) as evidenced by significantly higher blood velocity and patency compared to vehicle control beginning 40 minutes after injury and becoming significantly improved at 56 minutes (Figure 6H and Figure S3A).

## Discussion

The standard of care for occlusive cardiovascular diseases using DES is plagued by in-stent restenosis and thrombogenic risks[7, 9]. By far, efforts towards replacement of DES have not led to long-term clinical success. Here we provide evidence for an alternative anti-restenotic drug target; in other words, a potential therapeutic paradigm targeting PERK that may circumvent the drawbacks of DES. Specifically, we found that treatment with a selective PERK inhibitor mitigated not only IH but also thrombosis in preclinical models. These highly desirable in vivo outcomes are rationalized by the in vitro dual beneficial effects of PERK inhibition on both SMCs and ECs, i.e., not only abrogation of SMC de-differentiation/proliferation but also rescue of EC dysfunction. By contrast, increasing reports demonstrate that the anti-proliferative drugs used on DES (rapamycin and paclitaxel) exacerbate EC dysfunction[7, 9, 38].

In the present study, PERK inhibition proved effective in containing IH/restenosis in two drastically different (peri- and endovascular) drug-delivery routes. While the former was in line with a prospective anti-restenotic method suited for open surgeries (e.g. coronary artery bypass grafting), the latter using a biomimetic nanoplatform was conceived for a stent-free paradigm to prevent post-angioplasty restenosis, both elaborated in our recent publications[13, 22]. Bypass surgery and angioplasty account for the majority of vasculature reconstructions. Of particular interest, via endovascular i.v. administration of the PERK inhibitor GSK2606414 carried in lesion-homing biomimetic nanoclusters, we were able to reduce IH by >70% with a drug dose ~800-fold lower than those used for direct i.p. or i.v. injection in different disease models[20, 25]. Consistently, in our recent study, endovascular delivery using this biomimetic nanoplatform reduced the anti-restenotic effective dose of an epigenetic modulator (JQ1) by 70-fold compared to delivery of naked drug[13]. Whereas early studies using PERK inhibitors have noted systemic toxicity[20, 25], this barrier in therapeutic translation may be resolvable using our lesion-targeting endovascular delivery modality since it allows for the use of lower doses.

Lower effective dosages lend strong support to a target (PERK)-specificity of the anti-restenotic efficacy of GSK2606414 observed herein. This is an important feature especially in view of the use of pharmacological approaches to investigate the role for PERK in neointima formation. Arguing against possible off-target effects, the GSK2606414 dose in biomimetic nanoclusters is much below that in published studies that demonstrated a potent inhibition of PERK in several GSK2606414-treated tissues[20, 25]. These studies have confirmed that in an in vivo concentration range of 50-150mg/kg b.i.d., GSK2606414 is a highly selective inhibitor of PERK over other ER stress response kinases. Chances of GSK2606414 off-target effects were also reduced in our second route of delivery considering that the drug applied outside of the artery was not expected to enter circulation directly and quickly [26]. Furthermore, we observed a 4-fold dramatic upregulation of activated PERK in injured *vs* uninjured arteries at day 3 and day 7 post angioplasty, which should have substantially enhanced its targetability by GSK2606414. Strongly supporting a PERK target specificity, transgenically elevating PERK in the injured artery wall significantly enhanced IH. Taken together, these in vivo data of loss-of-function (PERK inhibitor) and gain-of-function (PERK up-regulation and overexpression) provide compelling evidence in support of a positive role for PERK in neointima development.

PERK’s pro-IH function is also supported by our in vitro results from loss-/gain-of-function experiments. Indicating a specific role of PERK, its silencing with an siRNA and overexpression with an adenoviral vector markedly inhibited and enhanced SMC phenotypic switching, respectively, as measured by cell proliferation and αSMA levels. It is well documented that SMC phenotypic switching is key to the development of IH[3, 4]. Furthermore, revealing the underlying PERK-mediated molecular mechanisms, PERK silencing and overexpression respectively reduced and increased the CDK1 protein, a cell division factor key to SMC proliferation. While a pro-proliferative role of PERK has been observed in cell types other than SMCs such as cancer cells[39], PERK-specific inhibition of αSMA expression, the hallmark of a differentiated SMC state, has not been previously reported. More detailed investigation uncovered that the PERK protein complexed with STAT3, a transcription factor known to promote SMC de-differentiation[33], and also MRTF-A which activates SRF, the master transcription factor governing αSMA expression and SMC differentiation[40]. The simplest interpretation of these results would be that the STAT3 protein activated or stabilized by the PERK kinase may sequester MRTF-A and inhibit its SRF-activating function.

Another important novel finding from this study is that PERK exacerbates EC dysfunction, particularly regarding the increased expression of tissue factor, a well-known potent thrombogenic factor[10]. While treatment with TNFα inhibited human aortic primary EC proliferation and dramatically up-regulated tissue factor protein by 40 fold, pretreatment with GSK2606414 abrogated this effect. Coincident with this result, in our parallel study of high throughput screening for drugs that inhibit SMC proliferation with lesser impact on EC viability, three PERK inhibitors including GSK2606414 and its derivatives were among the top hits (Figure S4C). Additional data in the current study indicated that the observed PERK function was probably mediated by NFκB (p65), the master transcription factor in ECs known to direct the expression of tissue factor[41]. Furthermore, the inhibitory effect of GSK2606414 on tissue factor expression was replicated in rat aorta samples in ex vivo culture, implicating an anti-thrombotic effect in vivo. Indeed, our in vivo experiments using the FeCl_3_-induced thrombosis model demonstrated that pre-treatment with GSK2606414 completely prevented arterial occlusion in 10 out of 16 mice, whereas untreated animals incurred nearly complete flow blockage without 60 min after induction with FeCl_3_. Consistent with this result, histology data confirmed much less thrombus inside of the arteries of GSK2606414-treated (*vs* untreated) mice.

The finding of anti-thrombotic efficacy of the PERK inhibitor is intriguing and somewhat serendipitous especially taking into account the anti-restenotic effect of the same PERK inhibitor first observed in this study. All the available therapeutics in the clinic have proven to be damaging to the fragile endothelium, a fact primarily underlying the increase of late and very late thrombosis incidence observed in patients receiving these therapies with DES[42]. On the other hand, hitherto reports on molecular targets that enable an anti-thrombotic and anti-restenotic dual beneficial intervention have been rare. Early studies identified various agents that inhibit SMC but not EC proliferation[38]. However, in most cases, their specific molecular targets remained unknown or unproven. Several recent reports pointed to potential interventional targets, such as ADAMTS7, and CTP synthase 1, for differential inhibition of SMC (*vs* EC) proliferation[14, 43]. However, these studies were conducted in global knockout animals, and hence their SMC- and EC-specific roles in vivo remained unclear. Moreover, currently there are no potent and selective small molecule inhibitors available for these proteins, limiting their potential for interventional targeting. Of particular interest, it was recently discovered that PIK75, a preclinical selective inhibitor of PI3K/p110α, exhibited both anti-restenotic and endothelium-protective effects in mouse models[9]. The PIK75 effects were similar to that of overexpressing PI3K/p85α[9, 44], which is the inhibitory subunit keeping the catalytic subunit PI3K/p110α inactive. While encouraging, achieving an anti-restenotic/anti-thrombotic dual beneficial clinical outcome by targeting PI3K/p110α or PI3K/p85α begs on an extraordinary drug selectivity in vivo. This appears challenging, given total 4 classes of PI3K heterodimers, each formed combinatorially with one of multiple regulatory subunits and one of multiple catalytic subunits that are functionally diverse or opposite[45, 46].

While we are the first to provide evidence for the dual beneficial role of PERK inhibition in mitigating IH and thrombosis, other ER stress response pathways have been previously reported to be significant regulators of SMC pathophysiology in vascular diseases. For example, ATF4, one of the downstream effectors in the extended PERK pathway, was implicated in the development of restenosis, particularly in the acute phase[47]. Our data showed a relatively delayed peak expression of ATF4 post angioplasty, possibly relevant to a different experimental setting or signaling cascade. Interestingly, another major ER stress response pathway, IRE1/XBP1, was shown to be actively involved in SMC differentiation; its dysregulation led to SMC phenotypic switching and neointima formation[16]. Literature evidence also implicates an involvement of ER stress response (or related) pathways in atherosclerosis and aneurysm[48, 49]. In aggregate, other reports and our current study underscore the importance of these pathways in SMC and EC pathophysiology and associated vascular diseases. A major future task would be to delineate the specific roles of these and other potential targets such as P85α/p110α[9, 44] and CTP synthase 1[14], in order to device optimized interventions with synergistic anti-restenotic/anti-thrombotic effects and minimal toxicity.

## Conclusions

This study provides evidence for an interventional paradigm of PERK targeting to achieve dual inhibition of SMC and EC dysfunction. In vitro, specific PERK silencing blocks SMC de-differentiation and proliferation and also rescues impaired EC growth. In vivo, treatment with a PERK inhibitor mitigates both restenosis and thrombosis. Given that PERK inhibitors are in human testing for different disease conditions, PERK targeting coupled with local-delivery technologies such as lesion-homing nanoplatforms may potentially translate into next-generation anti-restenotic therapies with low or no thrombogenicity.

## Author contributions

B.W., L-W.G. and K.C.K. conceived and designed the study. B.W. and M.Z. designed and performed all the in vitro experiments. B.W. and G.U. performed all the rat balloon angioplasty and in vivo gain-of-function experiments. D.W., D.D., A.H., and S.N. performed the mouse thrombosis studies and analyzed these data. G.C. and S.G. synthesized the biomimetic nanoclusters and perivascular tri-block gel for in vivo therapeutic experiments. Y.H. performed histology. B.W., M.Z., G.U., D.W., D.D., L-W.G., and K.C.K. analyzed the data. B.W. and L-W. G. co-wrote the manuscript.

## Acknowledgments

We thank GlaxoSmithKline for sharing the PKIS 2 library through the University of Wisconsin-Madison Small Molecule Screening Facility (SMSF). We thank Dr. Michael F Hoffmann for helpful discussion and technical support on drug screening, and we thank Song Guo and Gene Ananiev from SMSF) for their technical assistance. We thank Drew Alan Roenneburg’s technical support on histology.

## Sources of Funding

This work was supported by NIH grants R01HL143469, R01HL129785 (to K.C.K., S.G., L.-W.G.), R01HL133665 (to L.-W.G.), K25CA166178 (to S.G.), NCAI-CC Technology

Development Award (to S.N.), and AHA pre-doctoral awards 17PRE33670865 (to M.Z.) and 16PRE30160010 (to B.W.).

**Fig. S1.**
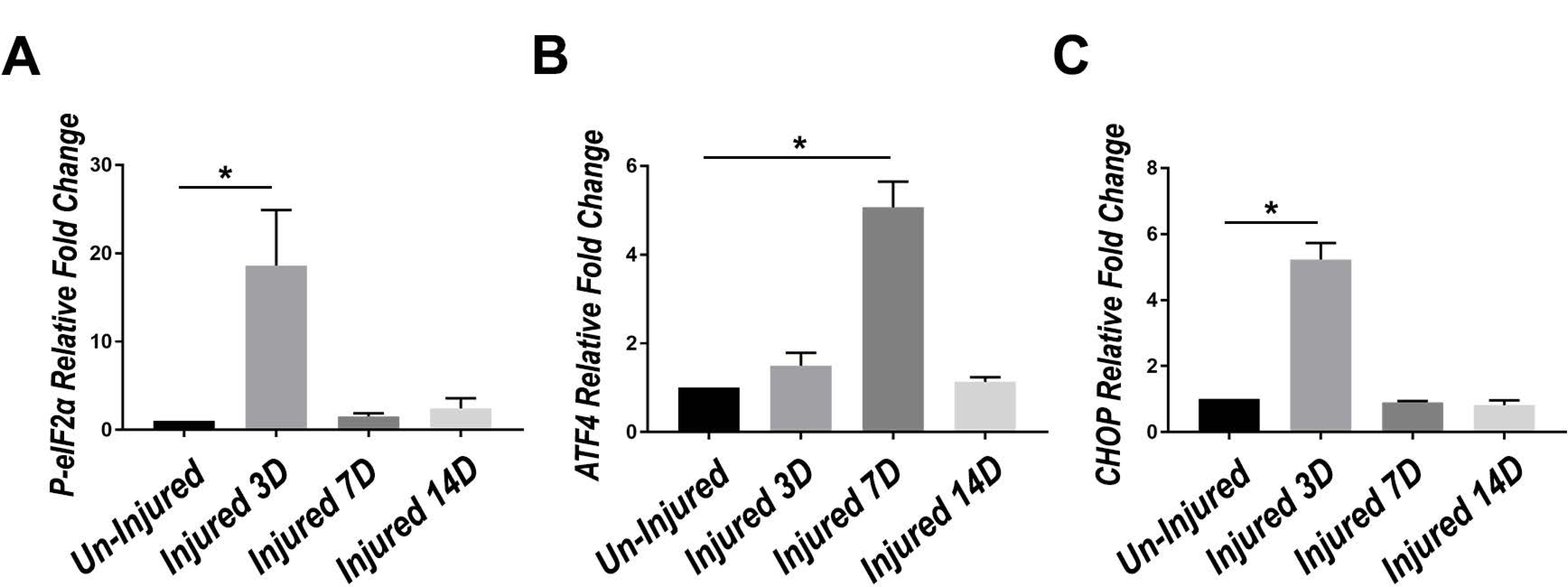
Quantitation of PERK pathway activation in balloon-injured rat carotid arteries. Quantitation of PERK pathway proteins in injured arteries following angioplasty as described in Fig.3. Rat common carotid arteries were harvested at days 3, 7, and 14 (3d-14d) and homogenized for immunoblotting analysis of P-eIF2α (A), ATF4 (B), and CHOP (C), respectively. Mean ± SEM, n=3 rats; *p<0.05, One-way ANOVA with Bonferroni post hoc test.

**Fig. S2.**
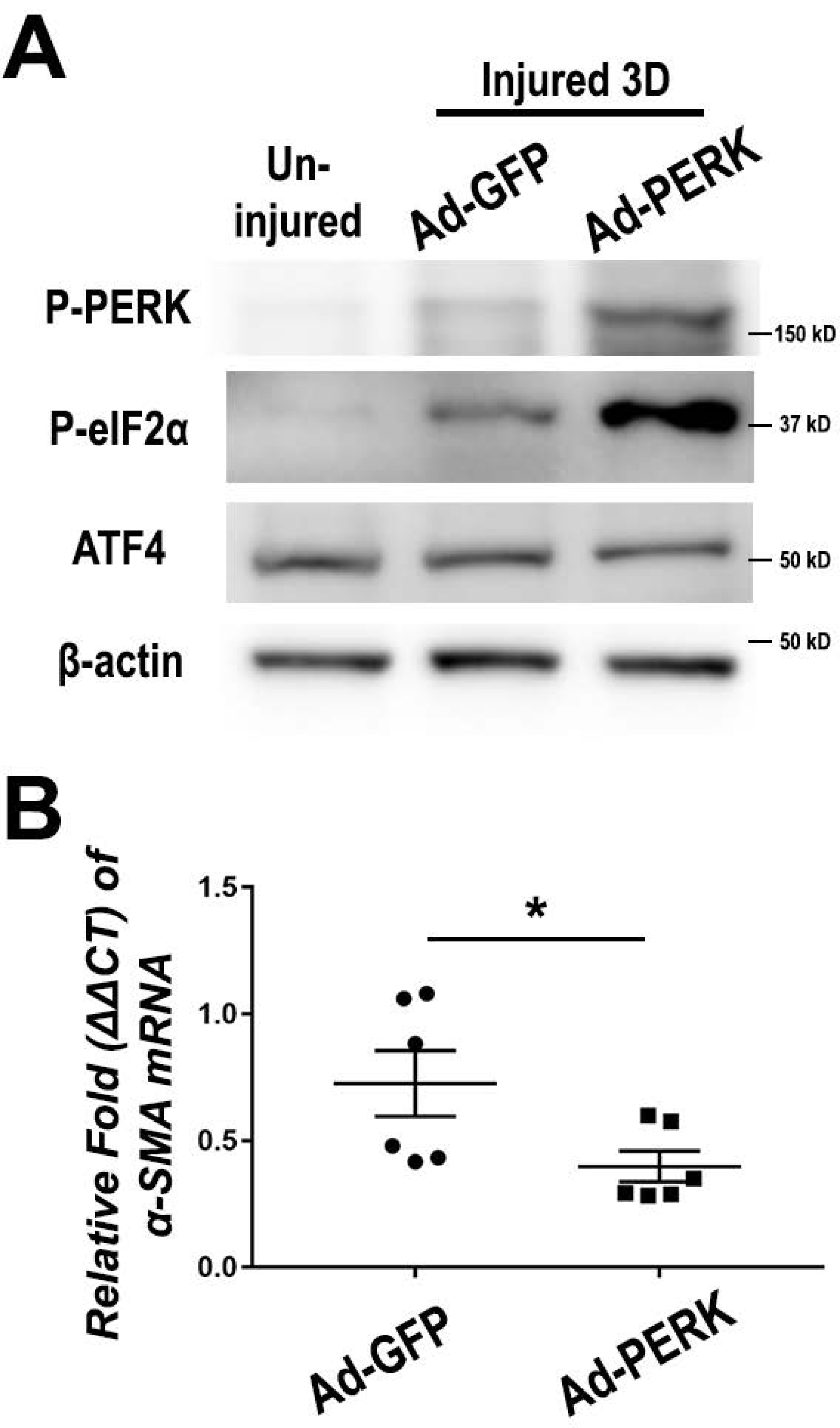
In vivo adenovirus-mediated PERK gain-of-function in balloon-injured rat carotid arteries. Rat carotid arteries were injured with balloon angioplasty, and subsequently locally infused with adenoviruses overexpressing GFP (Ad-GFP) or PERK (Ad-PERK) as described in method section. A. Adenovirus-infected rat carotid arteries were harvested at day 3 post injury for protein extraction and immunoblot analysis. In vivo infection efficiency was manifested by increased level of P-PERK and P-eIF2α. B. Adenovirus-infected rat carotid arteries were harvested at day 14 post injury for RNA extraction and qPCR analysis. Mean ± SEM, n=6 rats; *p<0.05, Mann-Whitney nonparametric test.

**Fig. S3.**
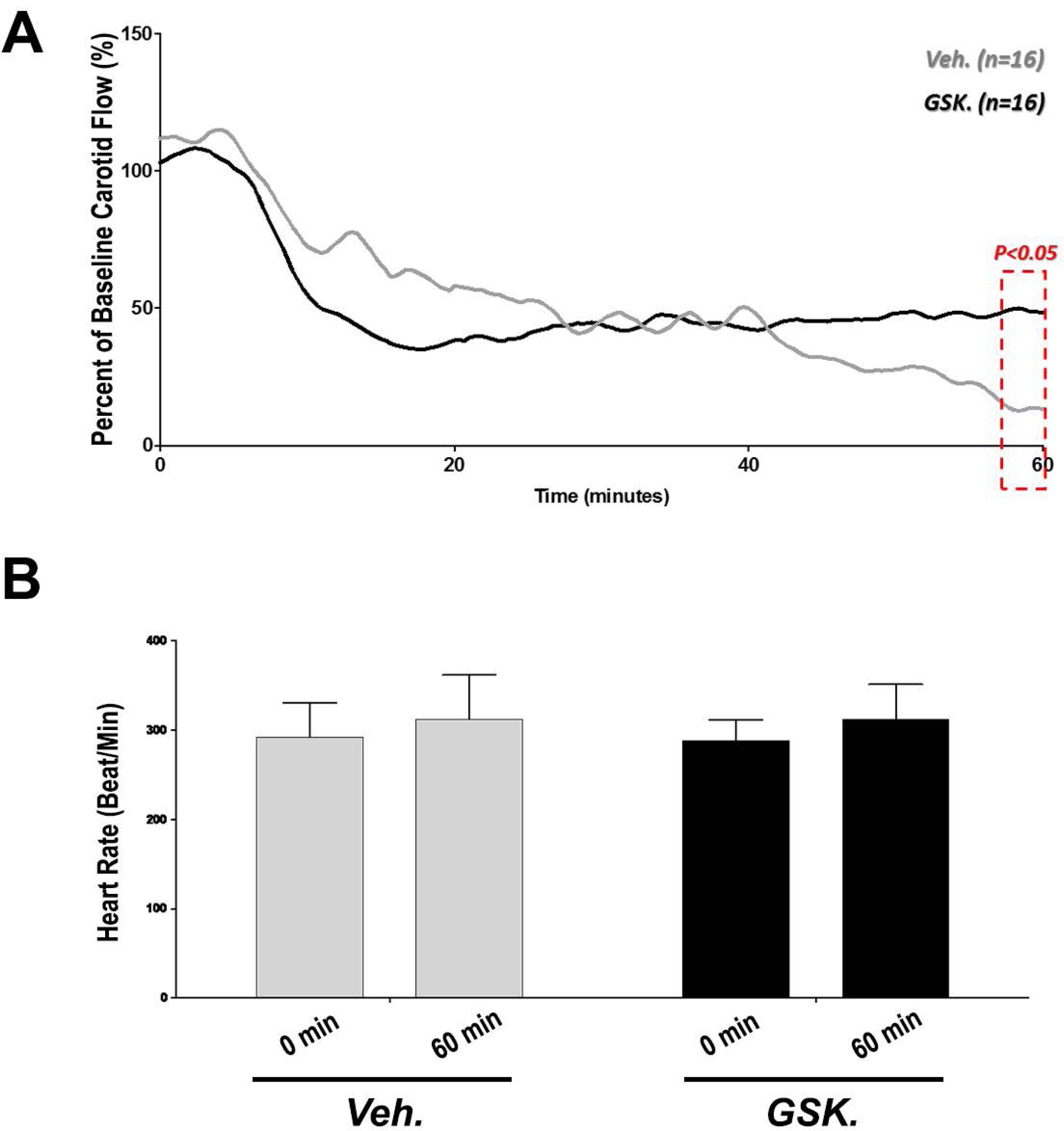
Effect of PERK inhibition in a murine FeCl_3_-induced thrombosis model. Mouse carotid arteries were locally applied with FeCl_3_-soaked patches to induce thrombosis formation. Mice were pre-treated with either vehicle or PERK inhibitor GSK2606414 (150 mg/kg) 4 hours prior to experiment. A. Real-time recording of blood velocity in FeCl_3_-injured carotid arteries. n=16 mice. B. Heart rate was monitored during the procedure. Initial and terminal heart rates are shown here. n=16 mice.

**Fig. S4.**
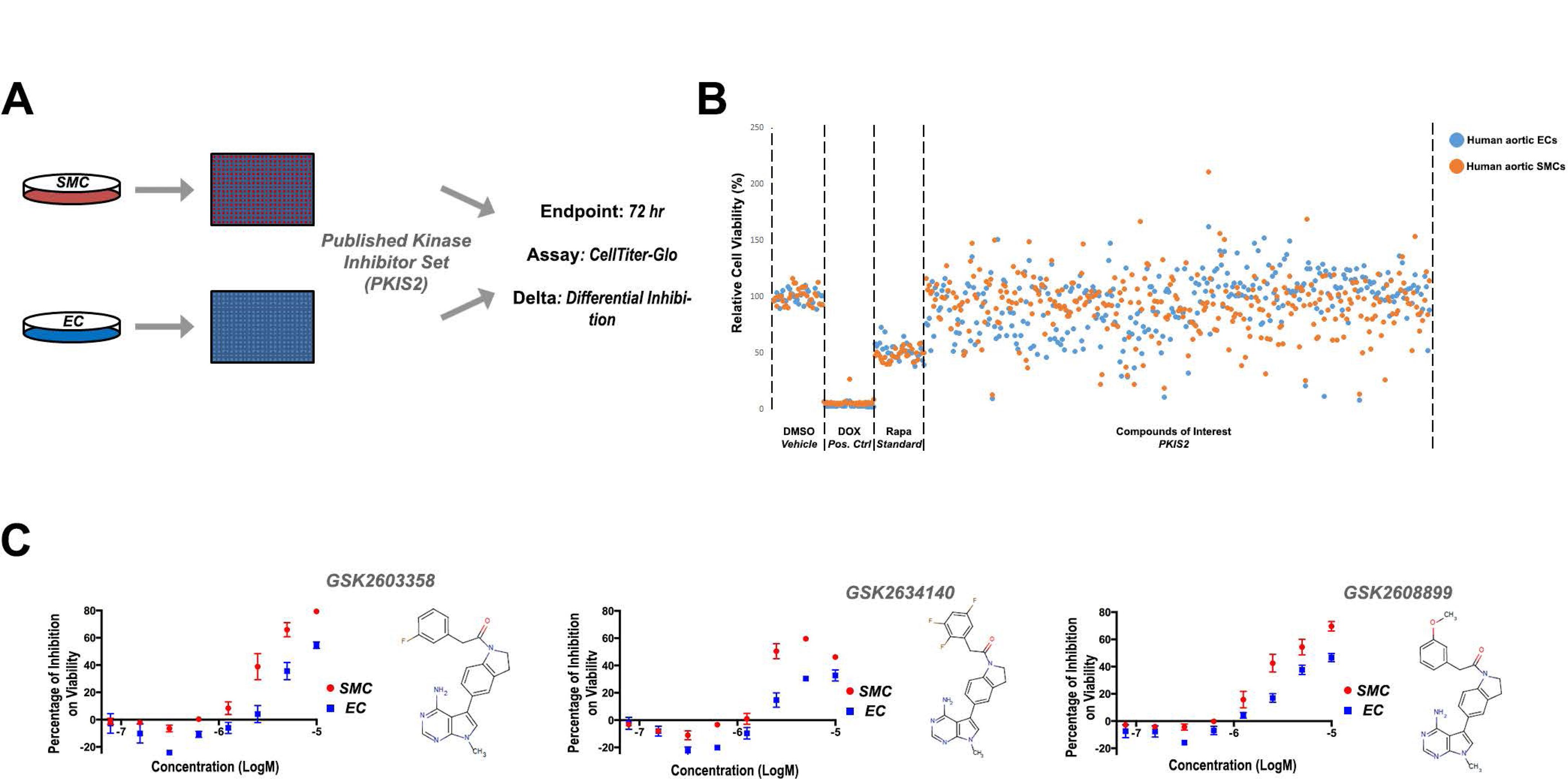
PERK inhibitors were identified with differential inhibition over SMC versus EC in a high-throughput phenotype screening. Human aortic SMCs and ECs were screened for their differential phenotypes to the PKIS 2 kinase inhibitor library as previously described. A. Scheme of the high-throughput drug screening. Cell viability was selected as the phenotype readout of the assay. B. Overview of the drug screening result. C. Among the identified hits with differential inhibition over SMC versus EC, we discovered 3 compounds that were previously shown with selective inhibition to PERK. These 3 compounds were derivatives of the first-in-class PERK inhibitor GSK2606414.

